# Extracellular Sulfatase Regulation of BMP-7 and FGF-2 Signaling in Human Articular Chondrocytes

**DOI:** 10.64898/2026.05.22.727020

**Authors:** Shuhei Otsuki, Shigeru Miyaki, Martin K. Lotz

**Affiliations:** Department of Orthopedic Surgery, Osaka Medical and Pharmaceutical University, Osaka Japan; Department of Histology and Molecular Cell Biology, Faculty of Medicine / Graduate School of Medicine, Kagawa University, Kagawa, Japan; Department of Molecular and Cellular Biology, The Scripps Research Institute, La Jolla, CA 92037

## Abstract

We performed a new set of experiments to confirm findings reported in our previous publication regarding the role of Sulf-1 and Sulf-2 in regulating BMP-7 and FGF-2 signaling in human articular chondrocytes. Using primary chondrocytes from five independent human donors, we examined the effects of Sulf knockdown on Smad1/5 and Erk1/2 phosphorylation.

Sulf-1 and Sulf-2 knockdown consistently reduced BMP-7–induced Smad1/5 phosphorylation and enhanced FGF-2–induced Erk1/2 phosphorylation. Although the magnitude of Erk1/2 activation was somewhat lower than originally reported, the direction and statistical significance of the effects were preserved.

These results confirm the original conclusions and support the role of Sulfs as dual regulators of BMP-7 and FGF-2 signaling pathways in human chondrocytes.

## Introduction

Sulf-1 and Sulf-2 are extracellular sulfatases that regulate heparan sulfate proteoglycan sulfation and modulate key signaling pathways, including BMP-7 and FGF-2 signaling (1,2).

In our previous study (3), we reported that Sulf knockdown decreases BMP-induced Smad activation while enhancing FGF-induced Erk activation in human chondrocytes. Questions were subsequently raised regarding specific figures.

To address these concerns, we performed independent experiments using newly prepared primary human chondrocytes, closely matching the original experimental design.

## Methods

### Chondrocyte isolation and siRNA transfection

Cartilage from human femoral condyles and tibial plateaus was obtained at autopsy from normal tissue donors and chondrocytes were isolated as described (3). Primary human articular chondrocytes were isolated from five independent donors with no history of joint disease.

Experiments were performed with high-density cultured chondrocytes in passage 1-2. Sulf expression was knocked down by 50nM siRNA (Sulf-1: s23298, Sulf-2: s31806, Thermo Fisher Scientific). Chondrocytes (0.35 × 10^6^) were seeded in 6 well plates, and transfection was performed using Lipofectamine RNAiMAX (Thermo Fisher Scientific). Cells were transfected with 50 nM siRNA for 48 h in medium containing 10% fetal bovine serum.

### Quantitative PCR (qPCR)

Gene expression was analyzed using TaqMan Gene Expression Assay probes for *Sulf-1* (Hs00392834_m1), *Sulf-2* (Hs01016480_m1), and *gapdh* (Hs02786624_g1) according to the manufacturer’s protocol (Thermo Fisher Scientific). Gene expression levels were assessed relative to GAPDH (n=5).

### Western blotting and densitometry

Human chondrocytes were stimulated with BMP-7 or FGF-2 (both at 100ng/ml, PeproTech) and cell lysates were prepared at the time points indicated. Sulf-1 and Sulf-2 proteins were determined by Western blotting using antibodies from Abcam (Sulf-1: ab172404, Sulf-2: ab232835) 1/1000, and peroxidase affinipure goat-anti Rabbit IgG (Jackson) 1/3000 in iBind Flex solution (Thermo Fisher Scientific). Western blotting was performed using lysates from five independent donors. Protein expression of phospho-Smad1/5, total Smad1, phospho-Erk1/2, and total Erk1/2 was analyzed using specific antibodies (Erk1/2: Santa Cruz Biotechnology; others: Cell Signaling Technology). The Western blots were analyzed by densitometry and band intensities were quantified using Image J and normalized to total protein. Densitometry results for each individual western blot experiment are shown in Suppl. Figures 2-4.

### Statistical analysis

Statistically significant differences were determined with a two-tailed Student’s t-test between two groups. The results are reported as mean ± SD. P < 0.05 was considered statistically significant.

## Results

### Efficient Sulf Knockdown

Human chondrocytes were transfected with siRNA targeting Sulf-1 or Sulf-2 and RNA was isolated 48h later for PCR analysis. Specific and significant reduction in Sulf mRNA expression was observed (n=5, *p< 0.01). siRNA treatment reduced Sulf-1 mRNA by 98% and protein by up to 84%, and Sulf-2 mRNA by 97% and protein by up to 43% (Fig. 1A, B). These reductions were consistent across independent donor samples (Suppl. Fig. 1A, B).

**Figure 1.**
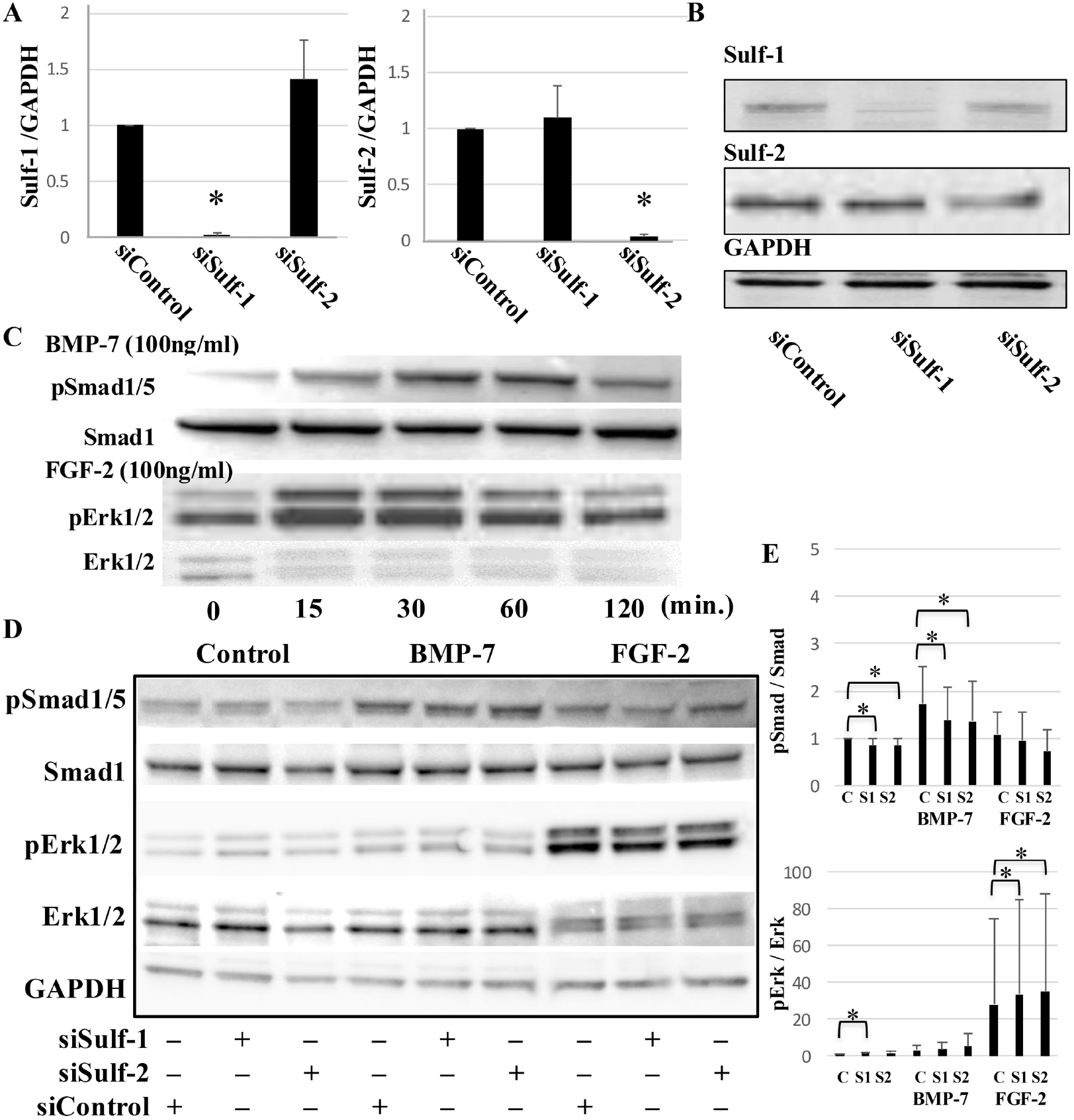
**(A)** Human chondrocytes were transfected with siRNA targeting Sulf-1 or Sulf-2 and RNA was isolated 48h later for PCR analysis. Specific and significant reduction in Sulf mRNA expression was observed (n=5, *p < 0.01). **(B)** Protein lysates were prepared 48 hours after siRNA transfection. Western blotting showed Sulf-1 and Sulf-2 protein levels were specifically reduced by 43-84 % following siRNA transfection (n=3-4). **(C)** Human chondrocytes were stimulated with BMP-7 or FGF-2 (both at 100 ng/ml), and cell lysates were collected at various time points as indicated (0–120 minutes) to examine Smad1/5 and Erk1/2 phosphorylation by Western blotting. In control cells, Smad1/5 and Erk1/2 phosphorylation increased in a time-dependent manner, peaking at 30–60 minutes after BMP-7 or FGF-2 stimulation, respectively (n=4). **(D)** Chondrocytes were transfected with Sulf siRNA to assess changes in Smad1/5 and Erk1/2 phosphorylation in response to BMP-7 and FGF-2 stimulation (100 ng/ml each). BMP-7–induced Smad1/5 phosphorylation was suppressed, whereas FGF-2–induced Erk1/2 phosphorylation was markedly enhanced in cells treated with Sulf-1 or Sulf-2 siRNA compared to control siRNA (n=5). The representative data were obtained from a 47-year-old male donor with normal cartilage. **(E)** Densitometry was performed on the Western blots from experiments with five different donors using NIH Image J. Results showed that BMP-7–induced Smad1/5 phosphorylation was significantly reduced in siSulf-1 and siSulf-2 treated cells (*p < 0.05, n=5). Conversely, siSulf-1 and siSulf-2 significantly enhanced FGF-2– induced Erk1/2 phosphorylation (*p < 0.05, n=5). Statistical analysis was performed using Student’s t-test between control and siRNA groups. C: siControl, S1: siSulf-1, S2: siSulf-2. *p<0.05.

### Time-Dependent Activation of BMP-7 and FGF-2 Signaling

In control cells, stimulation with BMP-7 or FGF-2 resulted in time-dependent increases in Smad1/5 and Erk1/2 phosphorylation, peaking at 30–60 minutes (Fig. 1C). Data for individual samples are shown in Suppl. Fig. 1C.

### Effects of Sulf Knockdown on BMP Signaling

Knockdown of Sulf-1 or Sulf-2 reduced basal Smad1/5 phosphorylation and significantly inhibited BMP-7– induced Smad1/5 phosphorylation (*p < 0.05) (Fig. 1D). These effects were consistent across all samples (Suppl. Fig. 1D).

### Effects of Sulf Knockdown on FGF Signaling

Sulf knockdown increased basal Erk1/2 phosphorylation (1.5–2 fold) and significantly enhanced FGF-2– induced Erk1/2 phosphorylation (Fig. 1D). Although the magnitude of Erk activation was somewhat lower than originally reported, the effect remained statistically significant (*p < 0.05, Fig. 1E). These effects were consistent across all donors (Suppl. Fig. 1D).

## Discussion

The present study independently confirms the key findings shown in Figure 6 of our original report (3). Sulf-1 and Sulf-2 knockdown consistently inhibited BMP-7–induced Smad activation while enhancing FGF-2– induced Erk activation in human chondrocytes. These findings were reproducible across multiple independent donor samples.

Although the magnitude of Erk1/2 activation was somewhat reduced compared to the original report, the direction and statistical significance of the effects were preserved. This variability may reflect donor-specific differences or differences in transfection efficiency.

Importantly, all experiments were repeated using newly prepared samples, and complete raw data are provided. These measures ensure transparency and reproducibility.

Taken together, these results support the conclusion that Sulfs act as dual regulators of BMP-7 and FGF-2 signaling pathways in human articular chondrocytes. Importantly, the direction of signaling regulation was consistent with the original findings across all independent experiments.

### Transparency Statement

All experiments were independently repeated using newly prepared primary chondrocytes from five donors. Full uncropped Western blot images and densitometry data are provided.

## Data Availability

All raw data and densitometry analyses are provided in Supplementary Figures.

## Notes

### Competing Interest Statement

The authors have declared no competing interest.

## References

1. Viviano BL, Paine-Saunders S, Gasiunas N, Gallagher J, Saunders S (2004) Domain-specific modification of heparan sulfate by Qsulf1 modulates the binding of the bone morphogenetic protein antagonist Noggin. J Biol Chem 279:5604–5611.

2. Wang S, Ai S, Freeman SD, Pownall ME, Lu Q, Kessler DS, Emerson JP (2004) QSulf1, a heparan sulfate 6-O-endosulfatase, inhibits fibroblast growth factor signaling in mesoderm induction and angiogenesis. Proc Natl Acad Sci USA 101:4833–4838.

3. Otsuki S, Hanson SR, Miyaki S, Grogan SP, Kinoshita M, Asahara H, Wong CH, Lotz MK (2010) Extracellular sulfatases support cartilage homeostasis by regulating BMP and FGF signaling pathways. Proc Natl Acad Sci USA 107:10202–10207.

