## Supplementary material for "Extracellular Sulfatase Regulation of BMP-7 and FGF-2 Signaling in Human Articular Chondrocytes": Suppl Figures

Suppl. Fig. 1A. Sulfs knock down with real time PCR

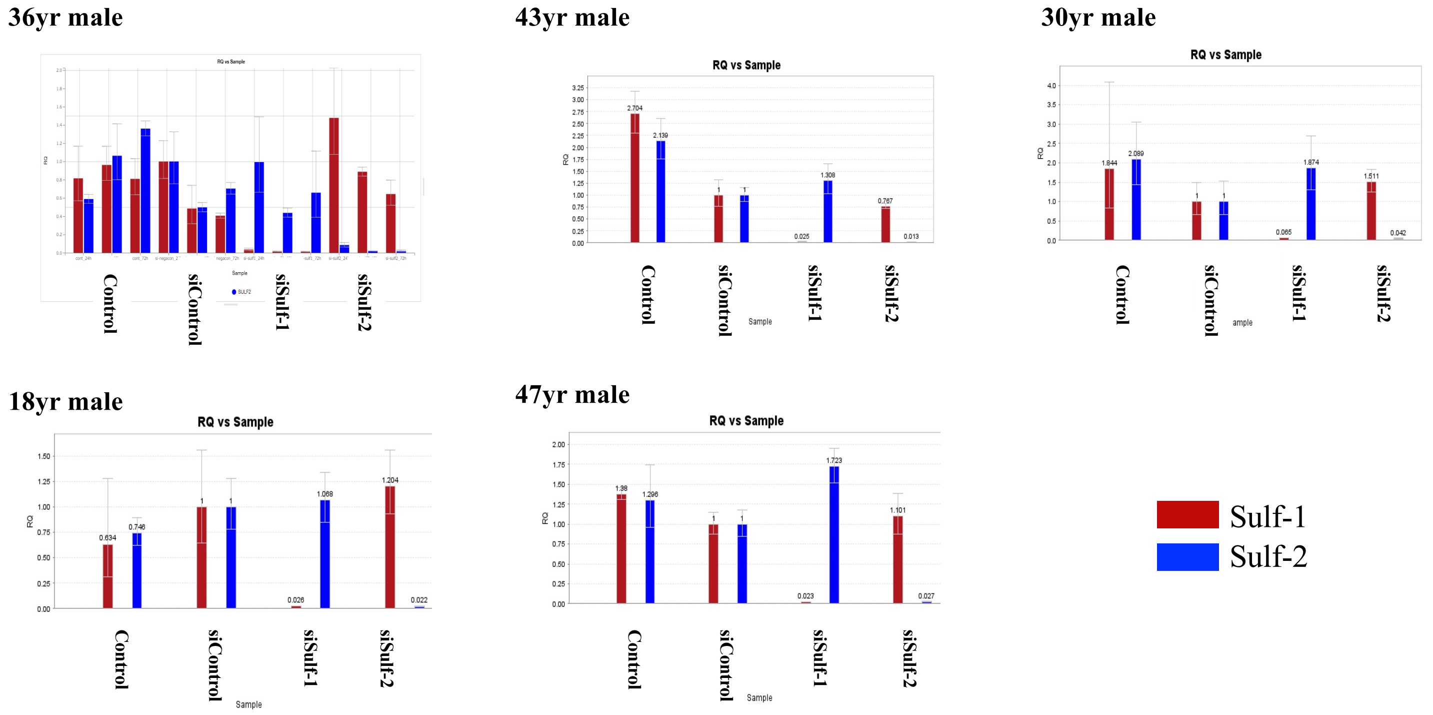

| Sulf-1 / GAPDH | 36yr M | 18yr M | 47yr M | 43yr M | 30yr M | Mean | SD |
|---|---|---|---|---|---|---|---|
| siCont. | 1 | 1 | 1 | 1 | 1 | 1 | 0 |
| siSulf-1 | 0.004 | 0.003 | 0.002 | 0.065 | 0.025 | 0.0198 | 0.027 |
| siSulf-2 | 1.07 | 1.09 | 1.72 | 1.84 | 1.31 | 1.41 | 0.36 |
| **Sulf-2 / GAPDH** | | | | | | | |
| siCont. | 1 | 1 | 1 | 1 | 1 | 1 | 0 |
| siSulf-1 | 0.92 | 1.2 | 1.1 | 1.5 | 0.77 | 1.1 | 0.28 |
| siSulf-2 | 0.06 | 0.02 | 0.03 | 0.042 | 0.013 | 0.033 | 0.019 |

#### Suppl. Fig. 1B: Sulfs knockdown with Western Blotting (Sulf-1)

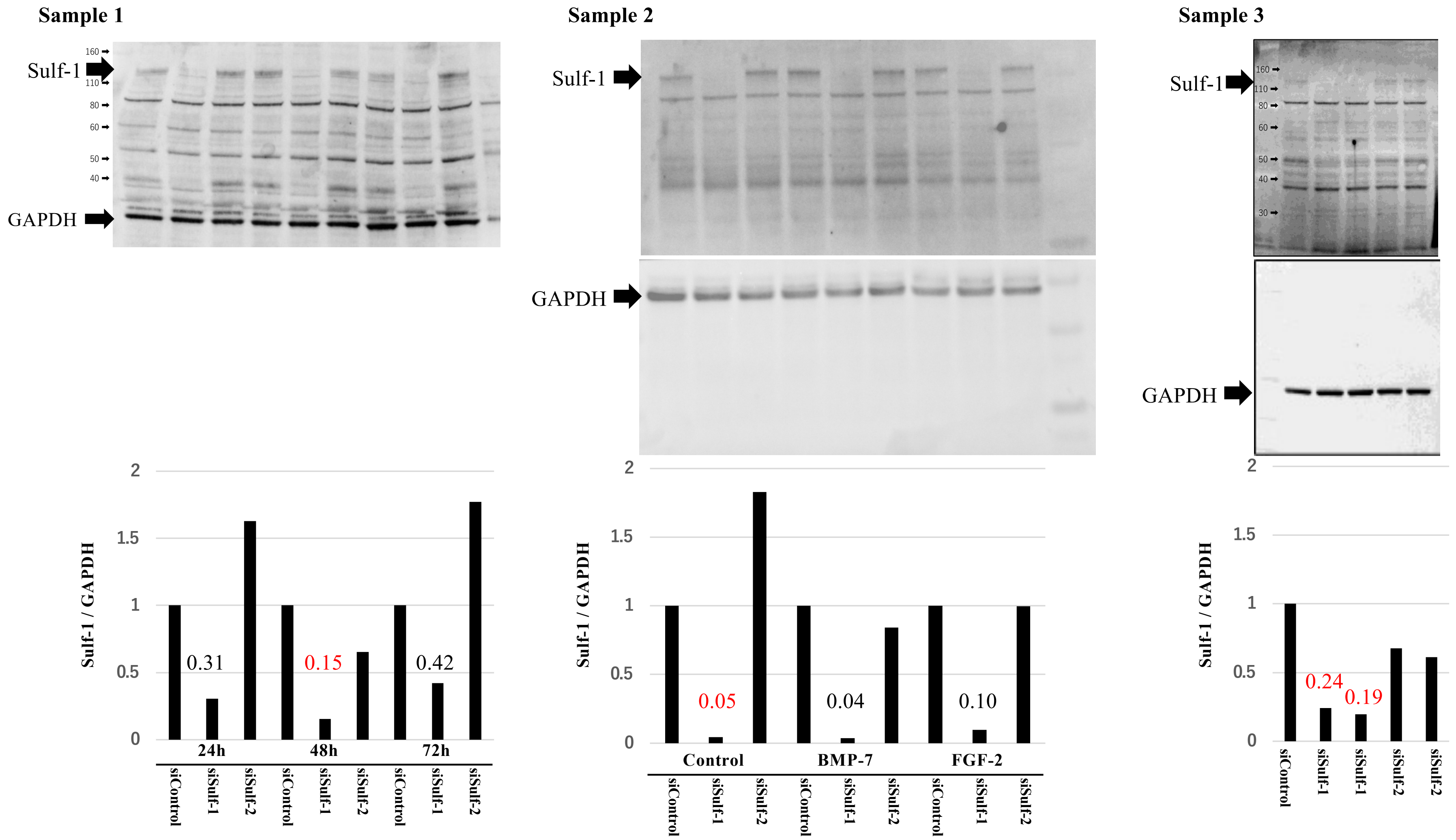

Sample: 20μg/lane (siControl, siSulf1, siSulf2) Transfected for 24h, 48h, and 72h, respectively
Sulf-1 (abcam, ab172404) Antibody 1/1000 in iBind Flex Solution
Peroxidase AffiniPure® Goat-anti Rabbit IgG (Jackson) 1/3000 in iBind Flex Solution

Suppl. Fig. 1B: Sulfs knockdown with Western Blotting (Sulf-2)

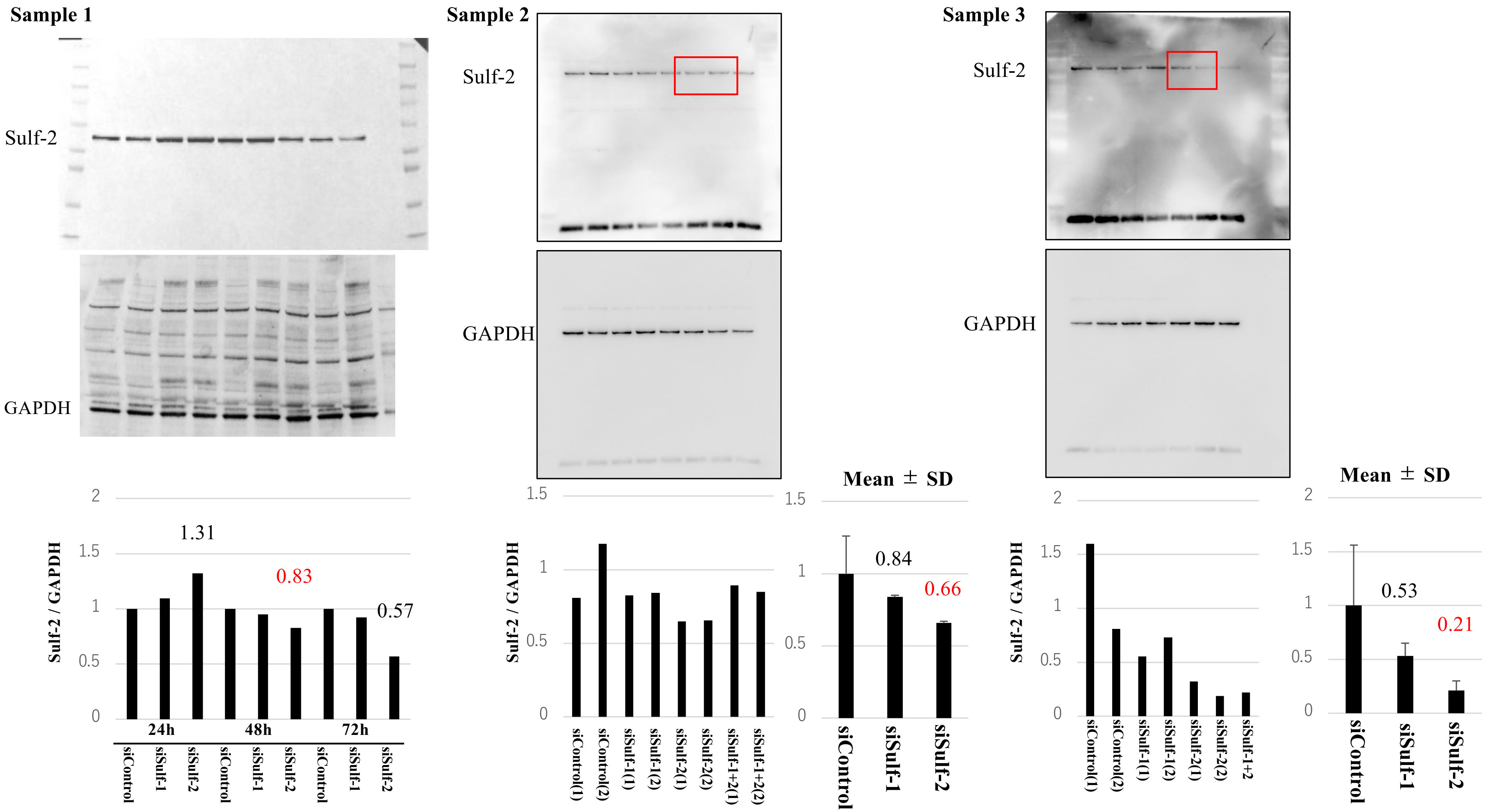

Sample: 20μg/lane
Sulf-2 (abcam, ab232835) Antibody 1/1000 in iBind Flex Solution
Peroxidase AffiniPure® Goat-anti Rabbit IgG (Jackson) 1/3000 in iBind Flex Solution
Duplicates with cells from same donors were examined. Sulf-1＋2 was the double knock down sample.

Suppl. Fig. 1C: Time course of pSmad1/5-Smad1 and pErk1/2-Erk1/2

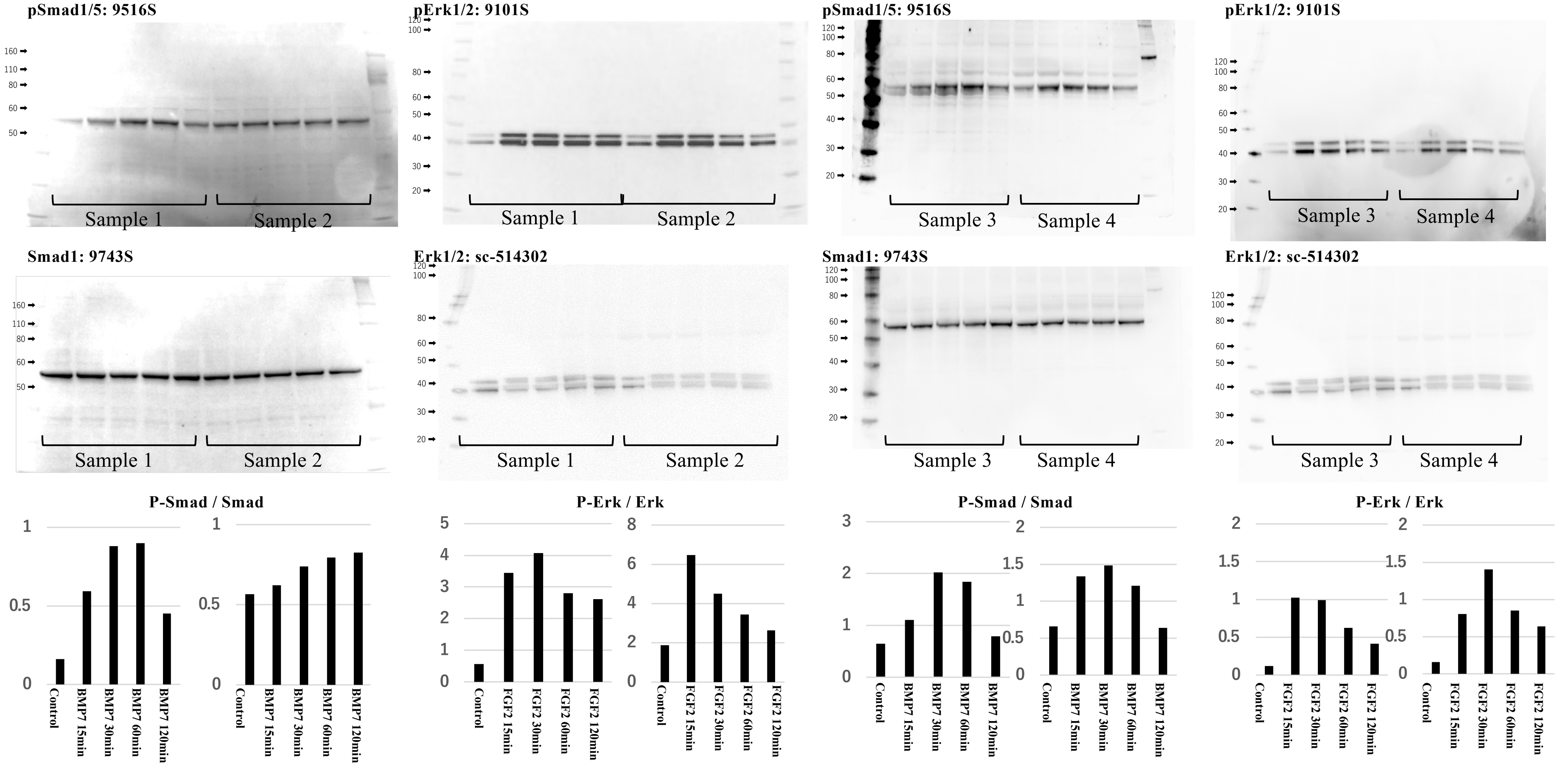

Sample: 20μg/lane
Phospho-SMAD1/5 (pSmad1/5) Antibody (cell signaling 9516S) 1/1000 in iBind Flex Solution. Peroxidase AffiniPure® Goat Anti-Rabbit IgG (Jackson) 1/3000 in iBind Flex Solution
Smad1 Antibody (cell signaling 9743S ) 1/1000 in iBind Flex Solution. Peroxidase AffiniPure® Goat Anti-Rabbit IgG (Jackson) 1/3000 in iBind Flex Solution
Phospho-p44/42 MAPK (pErk1/2) Antibody (cell signaling 9101S) 1/1000 in iBind Flex Solution. Peroxidase AffiniPure® Goat-anti Rabbit IgG (Jackson) 1/15000 in iBind Flex Solution
ERK 1/2 Antibody (Santa Cruz sc-514302) 1/1000 in iBind Flex Solution. Peroxidase AffiniPure® Goat Anti-Mouse IgG (Jackson) 1/15000 in iBind Flex Solution

Suppl. Fig. 1D: Sulfs knockdown in response to FGF2-Erk1/2 and BMP7-Smad1/5 signaling pathway

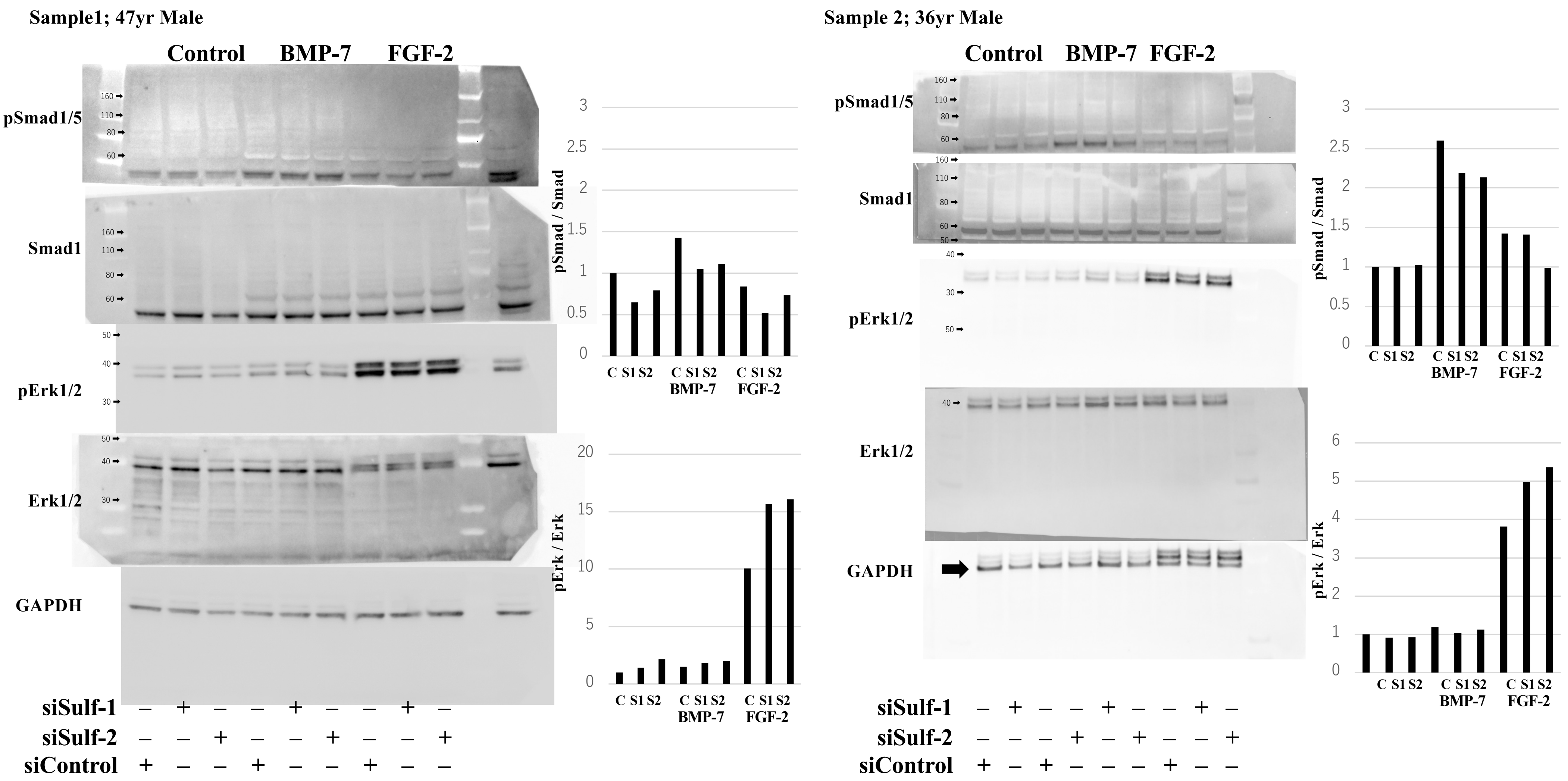

Suppl. Fig. 1D: Sulfs knockdown in response to FGF2-Erk1/2 and BMP7-Smad1/5 signaling pathway Continue

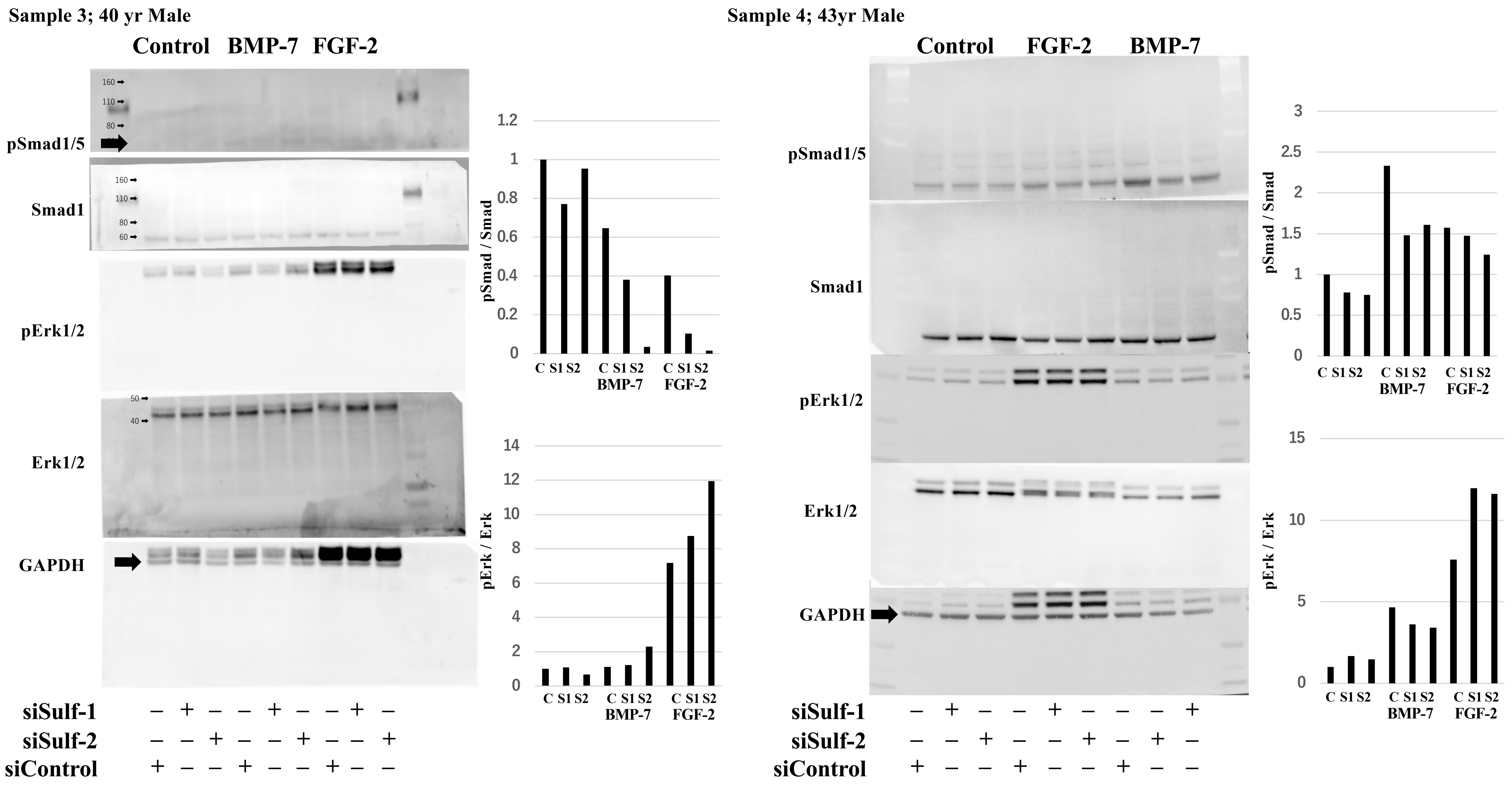

Suppl. Fig. 1D: Sulfs knockdown in response to FGF2-Erk1/2 and BMP7-Smad1/5 signaling pathway

Continue

**Sample 5; 46yr Male**

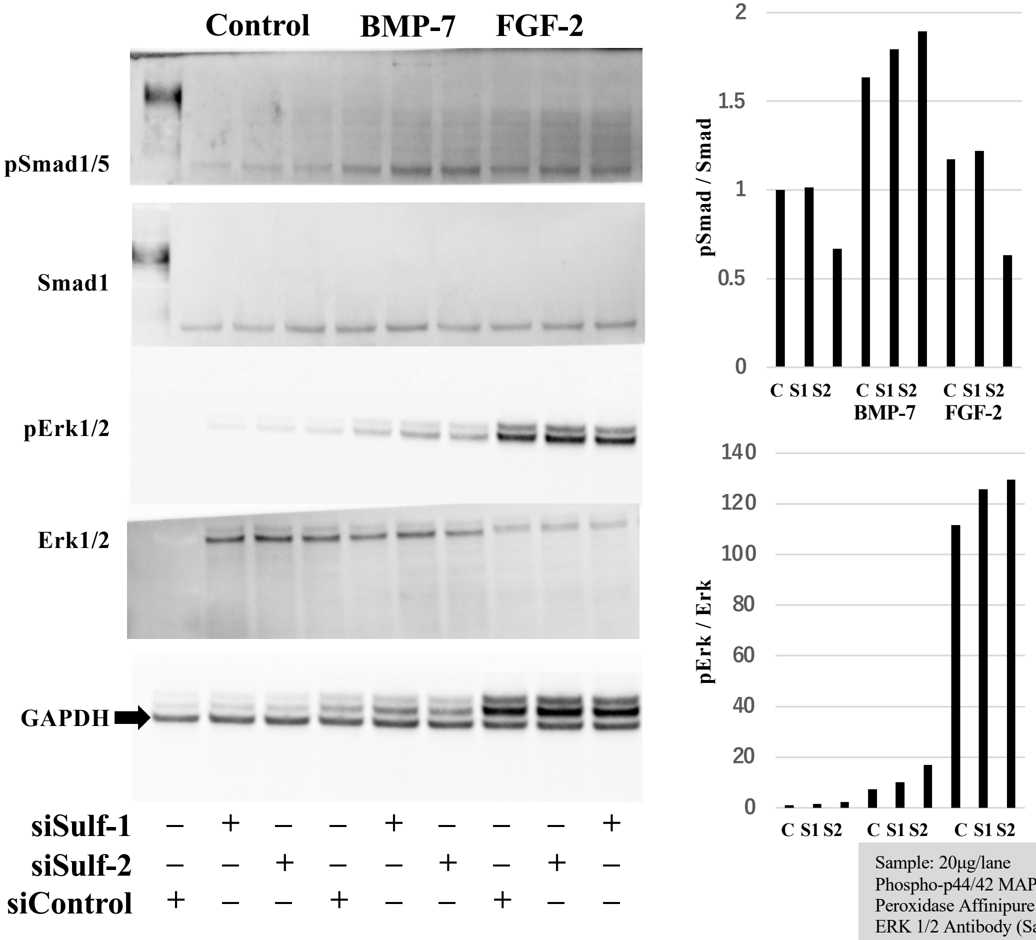

Sample: 20μg/lane
Phospho-p44/42 MAPK (pErk1/2) Antibody (cell signaling 9101S) 1/1000 in ibind Flex Solution
Peroxidase Affinipure Goat-anti Rabbit IgG (Jackson) 1/3000 in iBind Flex Solution
ERK 1/2 Antibody (Santa Cruz sc-514302 1/1000 in ibind Flex Solution, Peroxidase AffiniPure® Goat Anti-Mouse IgG (Jackson) 1/3000 in iBind Flex Solution
Phospho-Smad1/5 (pSmad1/5) Antibody (cell signaling 9516S) 1/1000 in ibind Flex Solution, Peroxidase AffiniPure® Goat Anti-Rabbit IgG (Jackson) 1/3000 in iBind Flex Solution
Smad1 Antibody (cell signaling 9743S ) 1/1000 in ibind Flex Solution, Peroxidase AffiniPure® Goat Anti-Rabbit IgG (Jackson) 1/3000 in iBind Flex Solution

### Suppl. Fig. 2A: Densitometry corresponding to Figure 1B (Sulf-1)

#### Sample 1

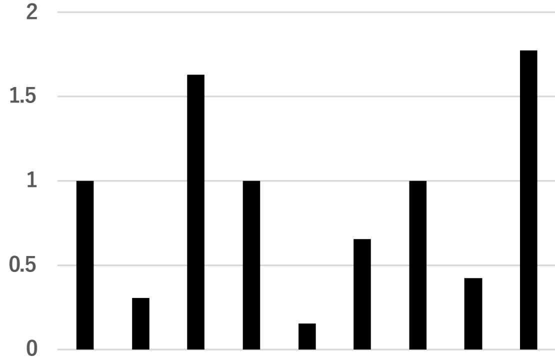

| | 24 hour | | | 48 hour | | | 72 hour | | |
|---|---|---|---|---|---|---|---|---|---|
| | Cont | siSulf-1 | siSulf-2 | Cont. | siSulf-1 | siSulf-2 | Cont. | siSulf-1 | siSulf-2 |
| Sulf-1 | 1283 | 310 | 2060 | 2247 | 381 | 1653 | 1160 | 438 | 2031 |
| GAPDH | 2036 | 1604 | 2009 | 162 | 1788 | 1830 | 1961 | 1754 | 1936 |
| Sulf-1 / GAPDH | 0.63 | 0.19 | 1.03 | 1.38 | 0.21 | 0.90 | 0.59 | 0.25 | 1.05 |
| % | 1 | 0.31 | 1.63 | 1 | 0.15 | 0.65 | 1 | 0.42 | 1.77 |

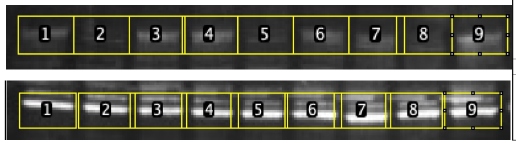

Sulf-1 GAPDH

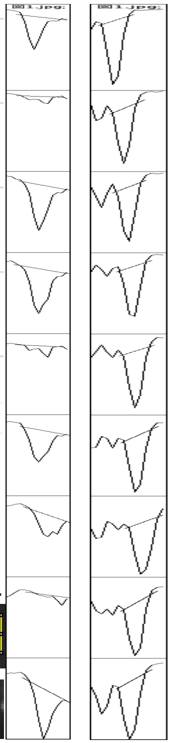

#### Sample 2

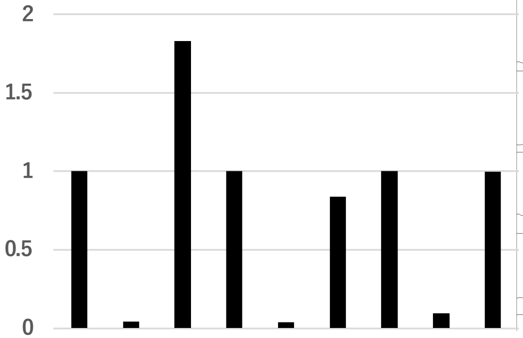

| | Control | | | BMP7 | | | FGF2 | | |
|---|---|---|---|---|---|---|---|---|---|
| | Cont | siSulf -1 | siSulf -2 | Cont. | siSulf -1 | siSulf -2 | Cont. | siSulf -1 | siSulf -2 |
| Sulf-1 | 8037 | 293 | 11753 | 15277 | 505 | 11042 | 8167 | 873 | 8546 |
| GAPDH | 24191 | 19335 | 19360 | 13108 | 11484 | 11269 | 11530 | 12572 | 12111 |
| Sulf-1 / GAPDH | 0.33 | 0.015 | 0.61 | 1.17 | 0.044 | 0.98 | 0.71 | 0.069 | 0.71 |
| % | 1 | 0.046 | 1.83 | 1 | 0.038 | 0.84 | 1 | 0.098 | 1.00 |

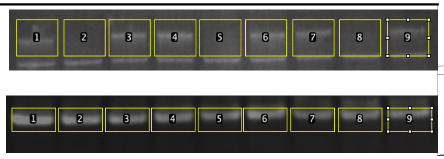

Sulf-1 GAPDH

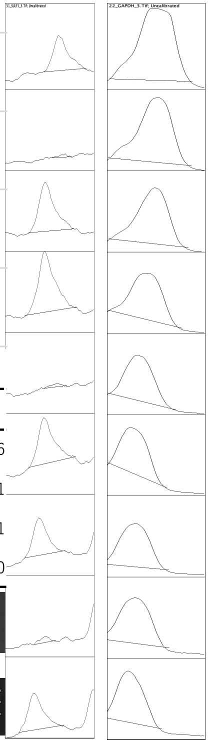

#### Sample 3

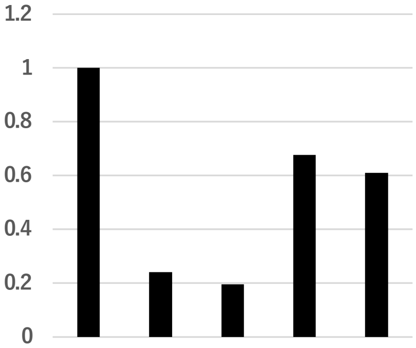

| | Cont | siSulf-1(1) | siSulf-1(2) | siSulf-2(1) | siSulf-2(2) |
|---|---|---|---|---|---|
| Sulf-1 | 8273 | 2438 | 1951 | 7531 | 6577 |
| GAPDH | 13710 | 16746 | 16604 | 18420 | 17832 |
| Sulf-1 / GAPDH | 0.60 | 0.15 | 0.12 | 0.41 | 0.37 |
| % | 1 | 0.24 | 0.19 | 0.68 | 0.61 |

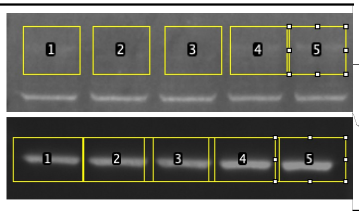

Sulf-1 GAPDH

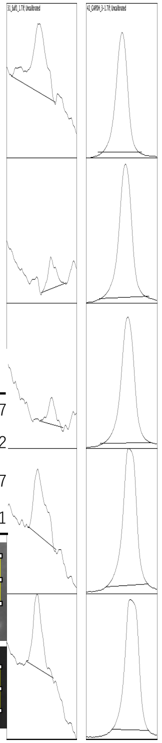

### Suppl. Fig. 2A: Densitometry corresponding to Figure 1B (Sulf-2)

#### Sample 1

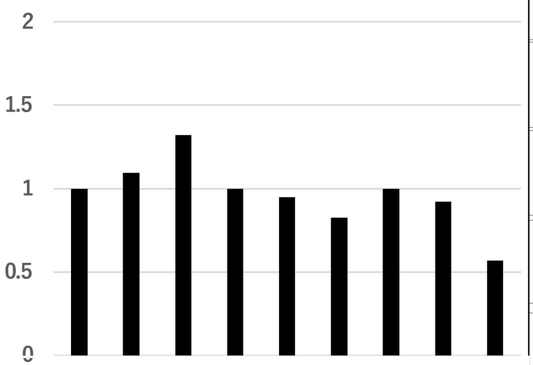

| | Cont | SiSulf-1 | SiSulf-2 | Cont. | SiSulf-1 | SiSulf-2 | Cont. | SiSulf-1 | SiSulf-2 |
|---|---|---|---|---|---|---|---|---|---|
| **Sulf-2** | 1801 | 1551 | 2348 | 2501 | 2608 | 2325 | 1708 | 1408 | 959 |
| **GAPDH** | 2036 | 1605 | 2009 | 1629 | 1788 | 1830 | 1961 | 1754 | 1936 |
| **Sulf-2 / GAPDH** | 0.88 | 0.97 | 1.17 | 1.53 | 1.46 | 1.27 | 0.87 | 0.80 | 0.50 |
| **%** | 1 | 1.09 | 1.32 | 1 | 0.95 | 0.83 | 1 | 0.92 | 0.57 |

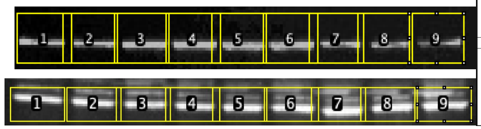

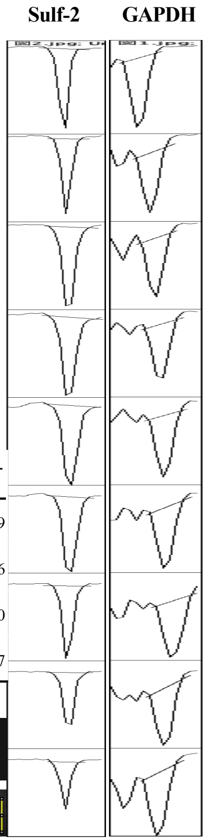

#### Sample 2

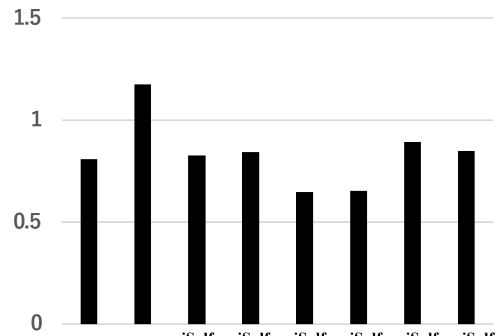

| | Cont.1 | Cont. 2 | siSulf-1(1) | siSulf-1(2) | siSulf-2(1) | siSulf-2(2) | siSulf-1+2(1) | siSulf-1+2(2) |
|---|---|---|---|---|---|---|---|---|
| **Sulf-2** | 170 | 1633 | 1239 | 1252 | 933 | 857 | 958 | 782 |
| **GAPDH** | 1696 | 1391 | 1501 | 1489 | 1443 | 1309 | 1073 | 923 |
| **Sulf-2 / GAPDH** | 0.81 | 1.17 | 0.83 | 0.84 | 0.65 | 0.66 | 0.89 | 0.85 |
| **Mean** | | 0.99 | | 0.83 | | 0.65 | | 0.87 |
| **SD** | | 0.26 | | 0.011 | | 0.0061 | | 0.03 |

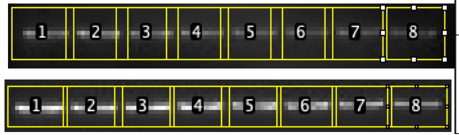

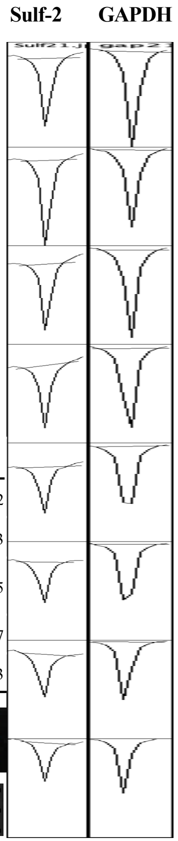

#### Sample 3

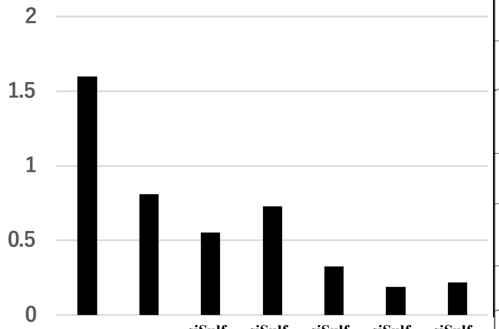

| | Cont. 1 | Cont. 2 | siSulf-1(1) | siSulf-1(2) | siSulf-2(1) | siSulf-2(2) | siSulf-1+2(1) |
|---|---|---|---|---|---|---|---|
| **Sulf-2** | 1324 | 942 | 809 | 1024 | 525 | 383 | 363 |
| **GAPDH** | 829 | 1161 | 1460 | 1403 | 1622 | 2021 | 1662 |
| **Sulf-2 / GAPDH** | 1.60 | 0.81 | 0.55 | 0.73 | 0.32 | 0.19 | 0.22 |
| **Mean** | | 1.21 | | 0.64 | | 0.26 | |
| **SD** | | 0.56 | | 0.12 | | 0.095 | |
| **%** | | 1 | | 0.53 | | 0.21 | |

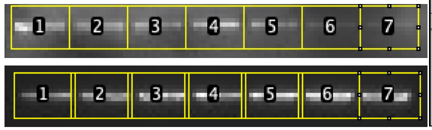

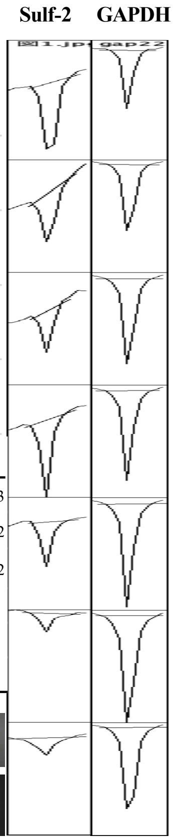

Suppl. Fig. 2B: Densitometry corresponding to Figure 1C (BMP-7 / Smad1/5 signaling)

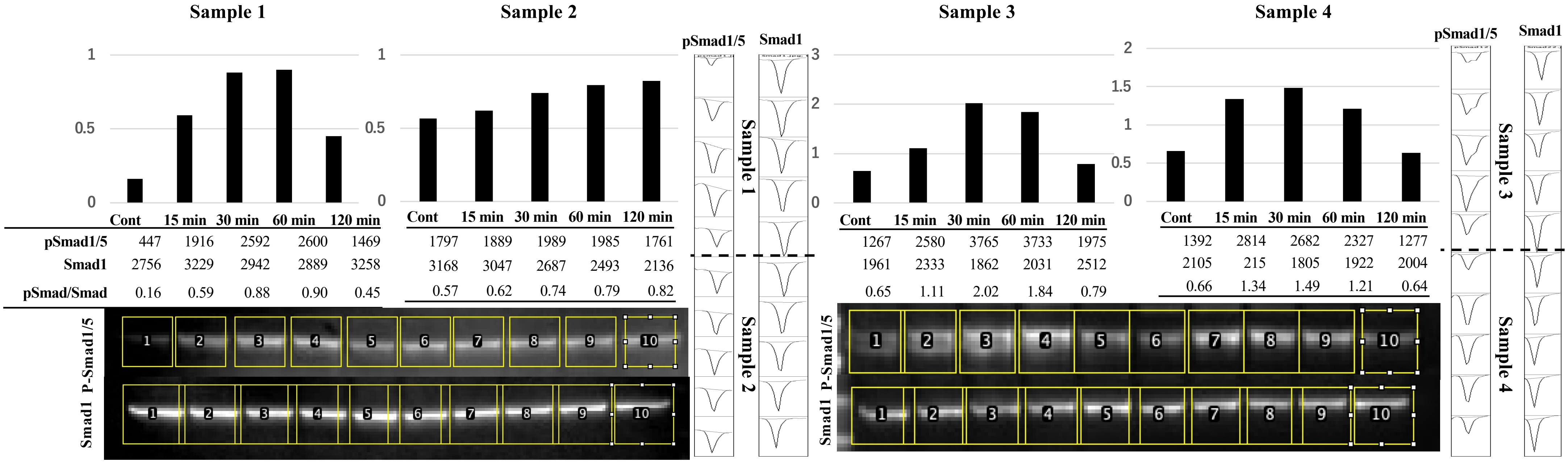

| | Sample | Cont | 15 min | 30 min | 60 min | 120 min |
|---|---|---|---|---|---|---|
| pSmad1/5 | Sample 1 | 447 | 1916 | 2592 | 2600 | 1469 |
| Smad1 | Sample 1 | 2756 | 3229 | 2942 | 2889 | 3258 |
| pSmad/Smad | Sample 1 | 0.16 | 0.59 | 0.88 | 0.90 | 0.45 |
| pSmad1/5 | Sample 2 | 1797 | 1889 | 1989 | 1985 | 1761 |
| Smad1 | Sample 2 | 3168 | 3047 | 2687 | 2493 | 2136 |
| pSmad/Smad | Sample 2 | 0.57 | 0.62 | 0.74 | 0.79 | 0.82 |
| pSmad1/5 | Sample 3 | 1267 | 2580 | 3765 | 3733 | 1975 |
| Smad1 | Sample 3 | 1961 | 2333 | 1862 | 2031 | 2512 |
| pSmad/Smad | Sample 3 | 0.65 | 1.11 | 2.02 | 1.84 | 0.79 |
| pSmad1/5 | Sample 4 | 1392 | 2814 | 2682 | 2327 | 1277 |
| Smad1 | Sample 4 | 2105 | 215 | 1805 | 1922 | 2004 |
| pSmad/Smad | Sample 4 | 0.66 | 1.34 | 1.49 | 1.21 | 0.64 |

Suppl. Fig. 2C: Densitometry corresponding to Figure 1C (FGF-2 / Erk1/2 signaling)

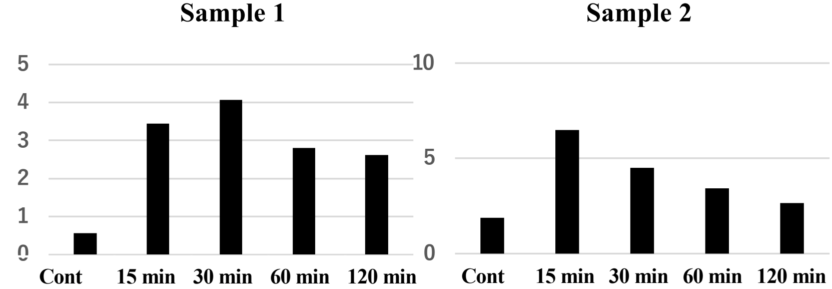

| | Sample 1: Cont | 15 min | 30 min | 60 min | 120 min | Sample 2: Cont | 15 min | 30 min | 60 min | 120 min |
|---|---|---|---|---|---|---|---|---|---|---|
| **pErk1/2** | 2144 | 10333 | 11197 | 10016 | 9650 | 5332 | 10366 | 10482 | 7852 | 5689 |
| **Erk1/2** | 3793 | 2996 | 2751 | 3565 | 3691 | 2849 | 1601 | 2322 | 2284 | 2150 |
| **pErk/Erk** | 0.57 | 3.45 | 4.07 | 2.81 | 2.61 | 1.87 | 6.47 | 4.51 | 3.44 | 2.65 |

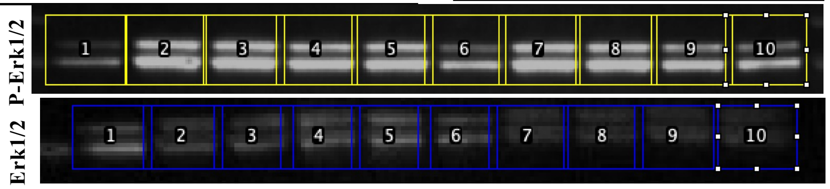

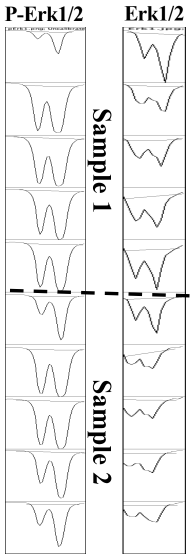

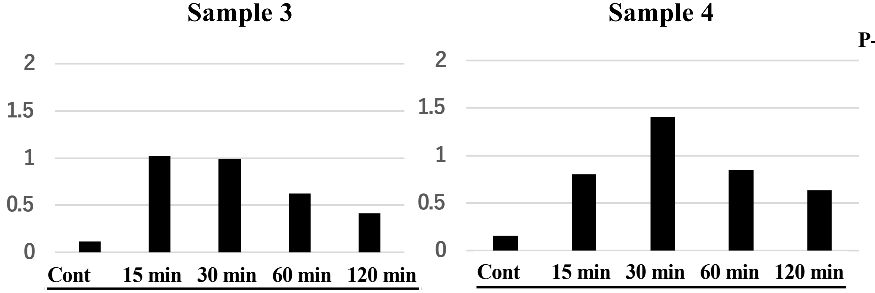

| | Sample 3: Cont | 15 min | 30 min | 60 min | 120 min | Sample 4: Cont | 15 min | 30 min | 60 min | 120 min |
|---|---|---|---|---|---|---|---|---|---|---|
| | 537 | 3764 | 3331 | 2701 | 1986 | 520 | 2021 | 2985 | 1735 | 1568 |
| | 4575 | 3672 | 3366 | 4312 | 4826 | 3322 | 2523 | 2123 | 2048 | 2468 |
| | 0.12 | 1.02 | 0.99 | 0.63 | 0.41 | 0.16 | 0.80 | 1.41 | 0.85 | 0.64 |

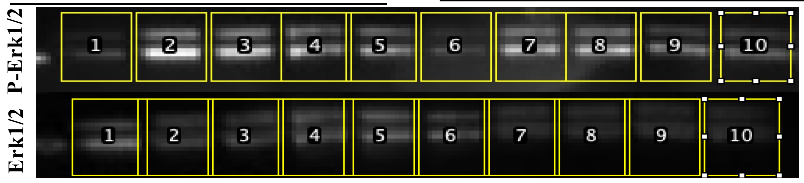

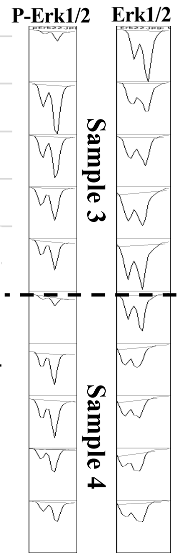

### Suppl. Fig. 3A: Densitometry corresponding to Figure 1D Sample 1

| | Control | | | BMP7 | | | FGF2 | | |
|---|---|---|---|---|---|---|---|---|---|
| | siCont | **siSulf-1** | **siSulf2** | **siCont** | **siSulf-1** | **siSulf2** | **siCont** | **siSulf-1** | **siSulf2** |
| pErk1/2 | 2999.953 | 5240.773 | 4095.51 | 5747.823 | 6965.359 | 7520.823 | 30023.798 | 28892.454 | 32188.354 |
| Erk1/2 | 9576.066 | 11695.187 | 6061.024 | 12338.978 | 12129.291 | 11905.907 | 9532.681 | 5894.882 | 6382.882 |
| pSmad1/5 | 8842.823 | 8129.338 | 6272.803 | 15227.986 | 12378.966 | 14066.501 | 8374.187 | 4543.196 | 6001.439 |
| Smad1 | 9393.066 | 13256.744 | 8424.53 | 11317.258 | 12498.329 | 13455.815 | 10610.016 | 9357.602 | 8647.874 |
| pErk/Erk | 0.31327614 | 0.44811366 | 0.67571255 | 0.46582651 | 0.57425937 | 0.63168837 | 3.14956495 | 4.90127775 | 5.04291854 |
| pSmad/Smad | 0.9414203 | 0.61322282 | 0.74458789 | 1.34555437 | 0.99044968 | 1.04538454 | 0.78927185 | 0.48550857 | 0.69397854 |
| **Ratio** | | | | | | | | | |
| **pErk/Erk** | 1 | 1.43041106 | 2.15692313 | 1.48695175 | 1.83307726 | 2.01639477 | 10.0536381 | 15.6452315 | 16.0973591 |
| **pSmad/Smad** | 1 | 0.6513805 | 0.79091973 | 1.42928124 | 1.05208023 | 1.1104334 | 0.83838415 | 0.51571925 | 0.73716123 |

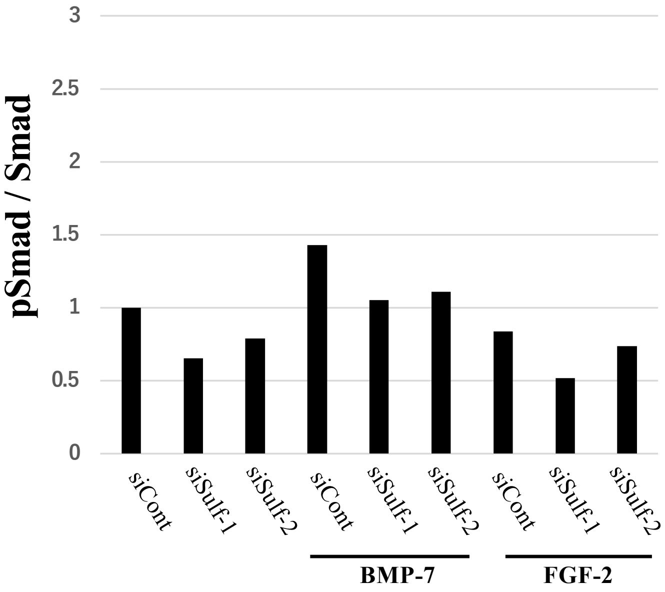

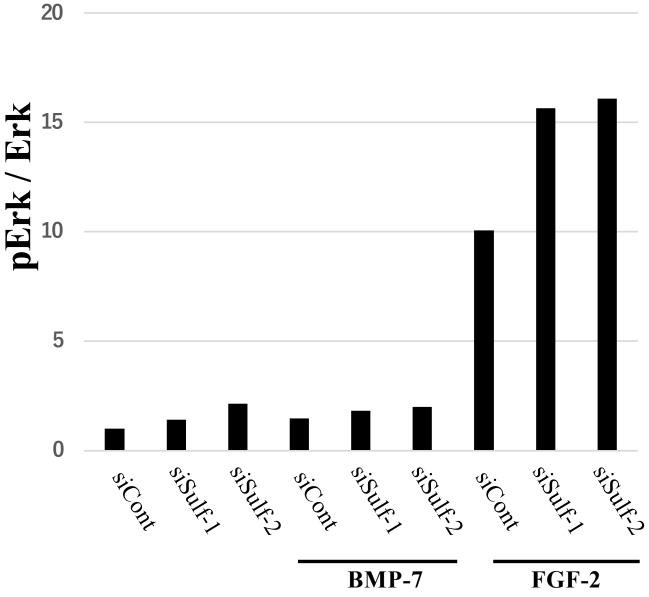

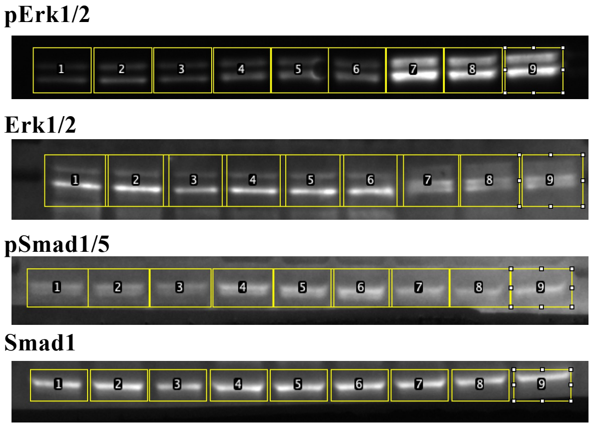

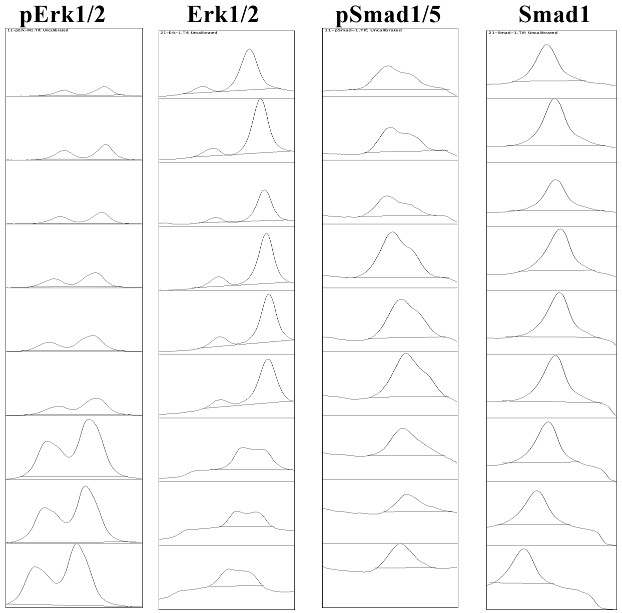

### Suppl. Fig. 3B: Densitometry corresponding to Figure 1D Sample 2

| | Control | | | BMP7 | | | FGF2 | | |
|---|---|---|---|---|---|---|---|---|---|
| | **siCont** | **siSulf-1** | **siSulf2** | **siCont** | **siSulf-1** | **siSulf2** | **siCont** | **siSulf-1** | **siSulf2** |
| pErk1/2 | 5563.832 | 3674.205 | 4483.104 | 5636.146 | 7549.116 | 6026.075 | 26205.413 | 23470.827 | 28461.555 |
| Erk1/2 | 21732.312 | 15717.785 | 18917.099 | 18462.149 | 28428.262 | 20970.827 | 26849.777 | 18405.442 | 20742.17 |
| pSmad1/5 | 9174.915 | 8162.238 | 8536.844 | 19596.907 | 19501.179 | 12188.158 | 7791.116 | 6030.681 | 5222.196 |
| Smad1 | 21300.108 | 18933.664 | 19309.472 | 17495.472 | 20686.472 | 13236.279 | 12729.744 | 9909.258 | 12239.652 |
| pErk/Erk | 0.25601657 | 0.23376099 | 0.23698687 | 0.30528115 | 0.26554968 | 0.28735514 | 0.97600114 | 1.27521127 | 1.37215899 |
| pSmad/Smad | 0.430745 | 0.43109659 | 0.44210655 | 1.12011308 | 0.94270202 | 0.92081453 | 0.61204027 | 0.60859057 | 0.42666213 |
| **Ratio** | | | | | | | | | |
| **pErk/Erk** | 1 | 0.91306977 | 0.92567002 | 1.19242729 | 1.03723631 | 1.12240837 | 3.81225769 | 4.98097159 | 5.35964911 |
| **pSmad/Smad** | 1 | 1.00081624 | 1.02637651 | 2.60040878 | 2.18853852 | 2.13772541 | 1.42088771 | 1.41287902 | 0.99052138 |

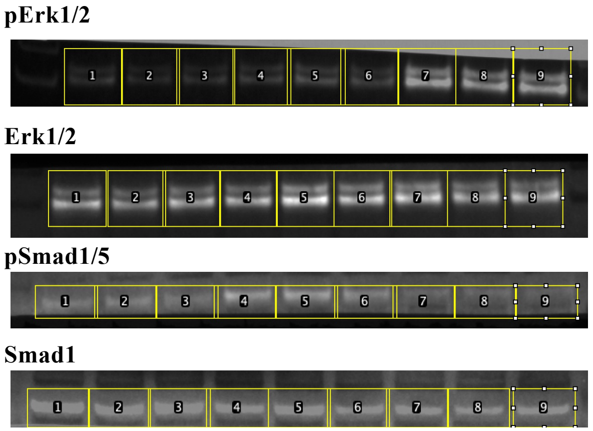

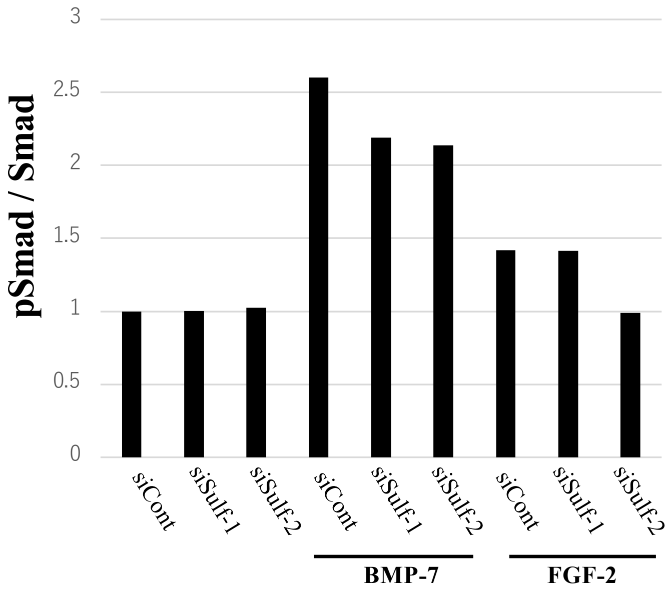

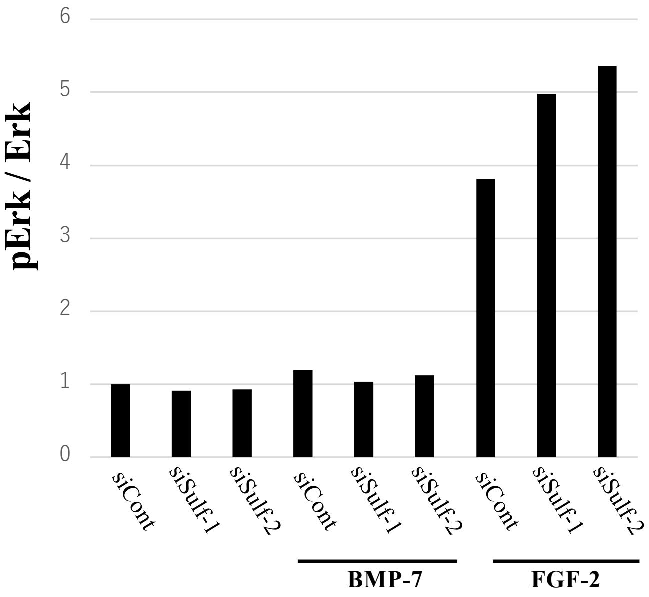

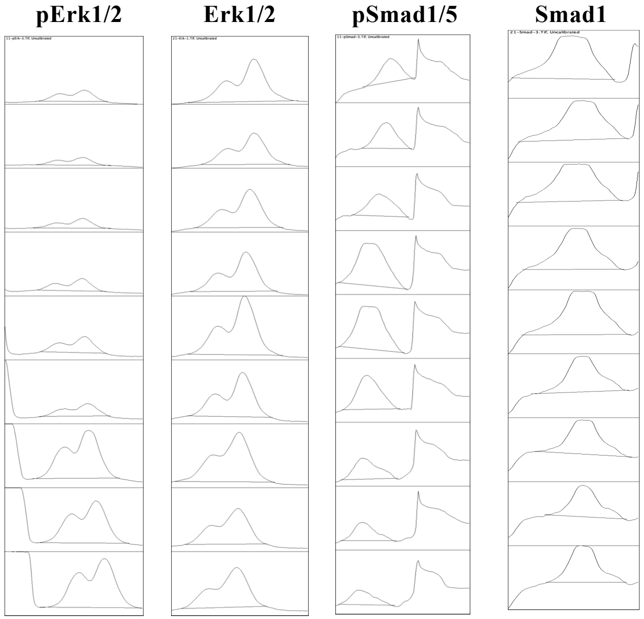

### Suppl. Fig. 3C: Densitometry corresponding to Figure 1D Sample 3

| | Control | | | BMP7 | | | FGF2 | | |
|---|---|---|---|---|---|---|---|---|---|
| | siCont | siSulf-1 | siSulf2 | siCont | siSulf-1 | siSulf2 | siCont | siSulf-1 | siSulf2 |
| pErk1/2 | 3390.891 | 4404.79 | 2355.749 | 5457.518 | 4307.305 | 9358.581 | 29129.321 | 28722.241 | 30112.534 |
| Erk1/2 | 16895.978 | 20072.463 | 17257.149 | 24310.605 | 17363.978 | 20258.685 | 20230.794 | 16334.522 | 12571.915 |
| pSmad1/5 | 11891.492 | 13903.484 | 14637.028 | 9912.409 | 4695.903 | 628.477 | 6664.731 | 1282.648 | 145.607 |
| Smad1 | 4838.024 | 7339.945 | 6236.51 | 6242.459 | 4997.267 | 7466.43 | 6736.995 | 5085.974 | 3695.66 |
| pErk/Erk | 0.2006922 | 0.21944442 | 0.13650859 | 0.22449125 | 0.24805981 | 0.46195402 | 1.43985061 | 1.75837658 | 2.39522253 |
| pSmad/Smad | 2.45792332 | 1.89422182 | 2.34699022 | 1.58790134 | 0.93969424 | 0.08417369 | 0.98927356 | 0.25219319 | 0.03939946 |
| **Ratio** | | | | | | | | | |
| **pErk/Erk** | 1 | 1.09343771 | 0.68018879 | 1.11858481 | 1.23602116 | 2.30180356 | 7.17442235 | 8.76155915 | 11.9348063 |
| **pSmad/Smad** | 1 | 0.77065945 | 0.95486715 | 0.64603372 | 0.38231227 | 0.03424586 | 0.40248349 | 0.10260417 | 0.01602957 |

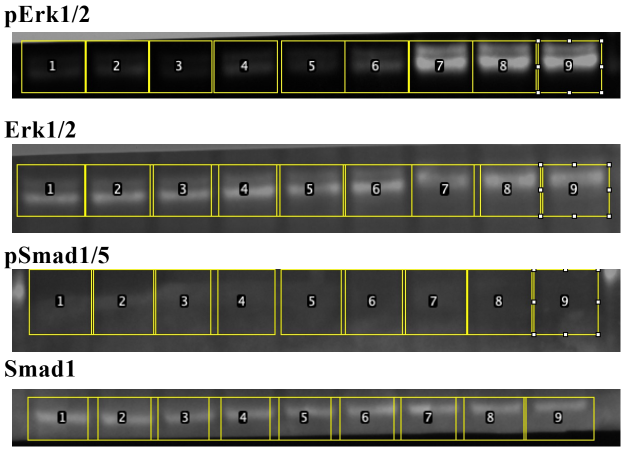

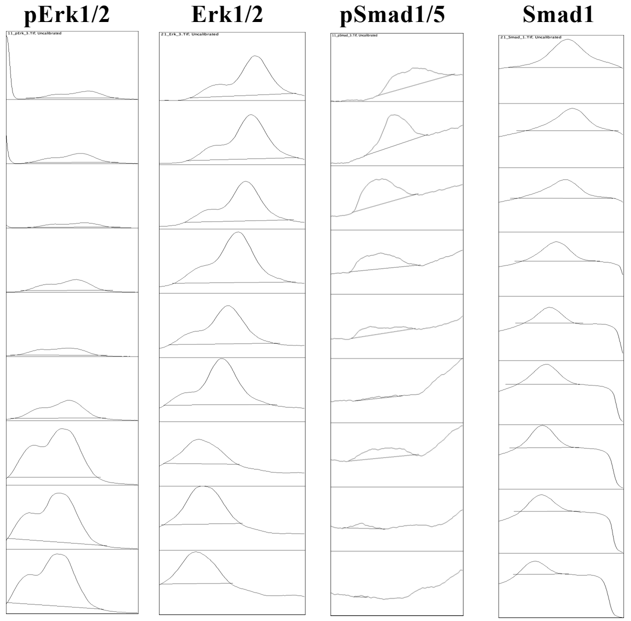

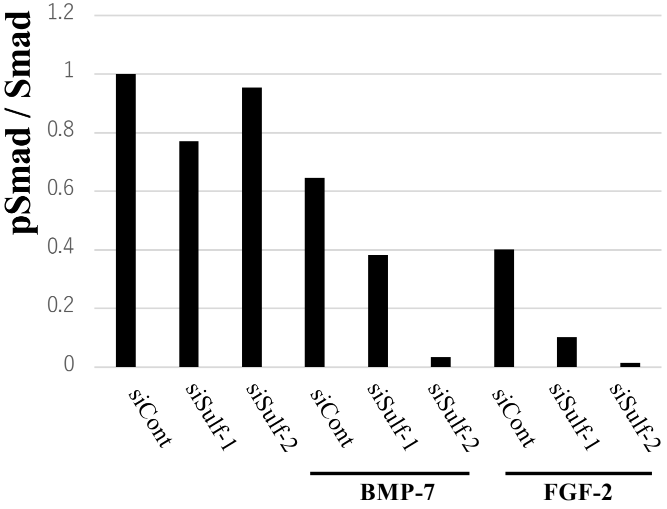

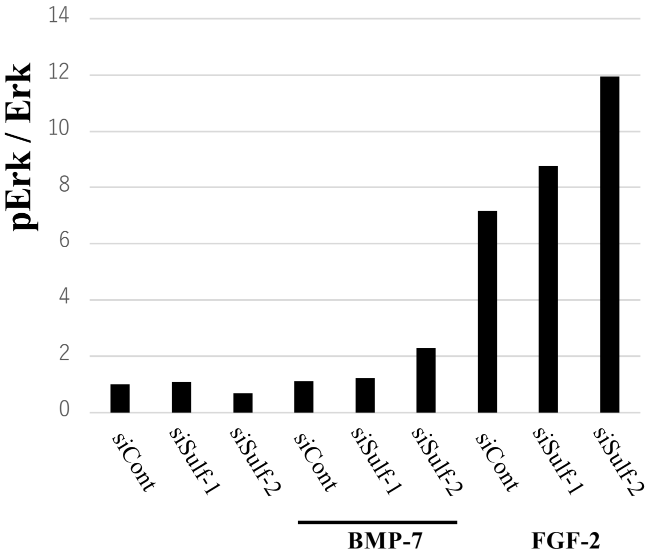

### Suppl. Fig. 3D: Densitometry corresponding to Figure 1D Sample 4

| | Control | | | BMP7 | | | FGF2 | | |
|---|---|---|---|---|---|---|---|---|---|
| | siCont | siSulf-1 | siSulf2 | siCont | siSulf-1 | siSulf2 | siCont | siSulf-1 | siSulf2 |
| P-Erk1/2 | 3965.317 | 6861.137 | 6151.551 | 9758.543 | 6637.773 | 9241.279 | 31122.304 | 34106.082 | 32547.082 |
| Erk1/2 | 18026.655 | 18564.241 | 19045.312 | 9535.057 | 8334.673 | 12279.886 | 18673.714 | 12976.087 | 12739.401 |
| P-Smad1/5 | 7747.602 | 7757.773 | 7940.602 | 24958.735 | 14060.2 | 14831.522 | 12763.836 | 11600.522 | 13208.057 |
| Smad1 | 10561.602 | 13601.208 | 14401.309 | 14602.501 | 12963.551 | 12590.966 | 11049.066 | 10710.066 | 14517.208 |
| P-Erk/Erk | 0.21996965 | 0.36958888 | 0.32299555 | 1.02343835 | 0.79640473 | 0.75255414 | 1.66663707 | 2.62837957 | 2.55483613 |
| P-Smad/Smad | 0.73356315 | 0.57037382 | 0.55138057 | 1.70920961 | 1.0845948 | 1.17794949 | 1.15519592 | 1.08314197 | 0.90982075 |
| **Ratio** | | | | | | | | | |
| **P-Erk/Erk** | 1 | 1.68018123 | 1.46836415 | 4.65263434 | 3.62052096 | 3.42117258 | 7.57666827 | 11.9488282 | 11.6144938 |
| **P-Smad/Smad** | 1 | 0.77753882 | 0.75164705 | 2.33001019 | 1.47852956 | 1.60579153 | 1.57477366 | 1.47654905 | 1.24027597 |

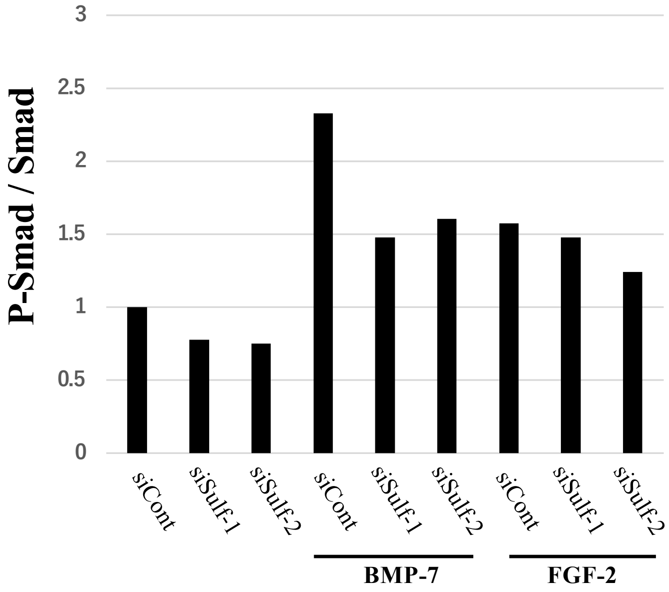

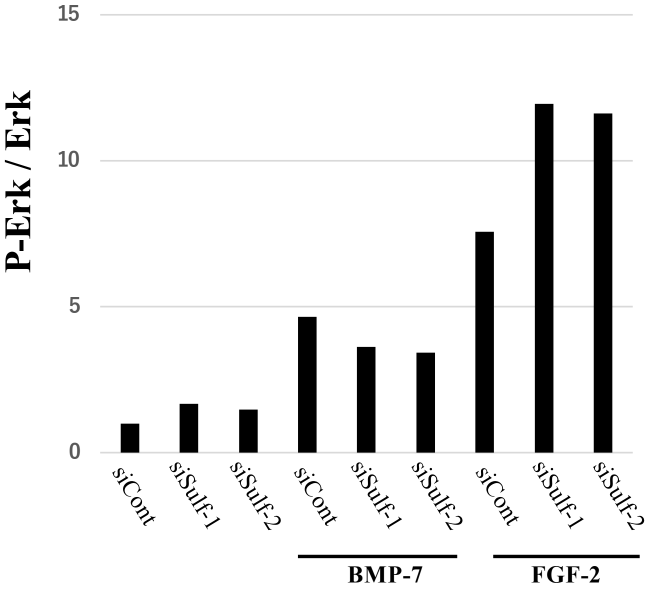

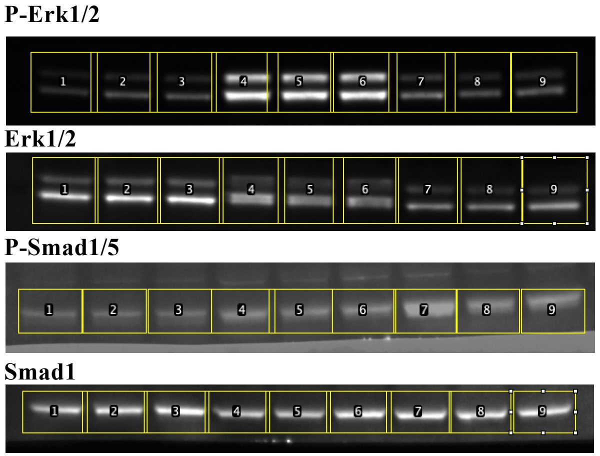

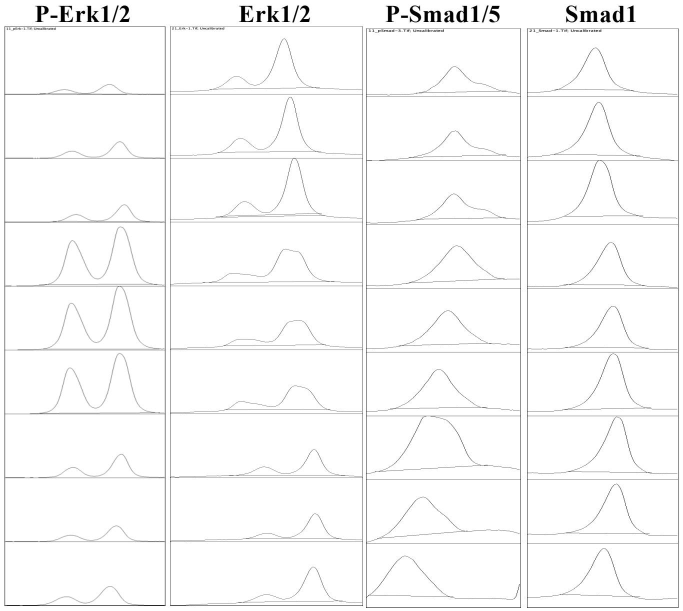

### Suppl. Fig. 3E: Densitometry corresponding to Figure 1D Sample 5

| | Control | | | BMP7 | | | FGF2 | | |
|---|---|---|---|---|---|---|---|---|---|
| | siCont | siSulf-1 | siSulf2 | siCont | siSulf-1 | siSulf2 | siCont | siSulf-1 | siSulf2 |
| P-Erk1/2 | 898.678 | 1641.163 | 2138.062 | 5653.903 | 9691.894 | 10813.38 | 33944.919 | 39542.768 | 30149.09 |
| Erk1/2 | 9461.238 | 11574.723 | 9979.48 | 8228.116 | 10015.723 | 6696.459 | 3198.861 | 3310.347 | 2452.548 |
| P-Smad1/5 | 1888.305 | 2226.033 | 2458.376 | 6520.459 | 8169.945 | 6746.581 | 3718.832 | 4566.196 | 5637.154 |
| Smad1 | 3427.882 | 3985.66 | 6693.045 | 7236.581 | 8275.409 | 6459.095 | 5755.217 | 6786.631 | 7578.773 |
| P-Erk/Erk | 0.09498524 | 0.14178853 | 0.21424583 | 0.6871443 | 0.96766794 | 1.61479074 | 10.6115642 | 11.9452033 | 12.2929663 |
| P-Smad/Smad | 0.5508664 | 0.55851051 | 0.36730307 | 0.90104139 | 0.98725574 | 1.04450871 | 0.64616712 | 0.6728222 | 0.74380827 |
| **Ratio** | | | | | | | | | |
| **P-Erk/Erk** | 1 | 1.49274274 | 2.25556964 | 7.23422155 | 10.1875607 | 17.0004379 | 111.718029 | 125.758516 | 129.419748 |
| **P-Smad/Smad** | 1 | 1.01387653 | 0.66677343 | 1.63568044 | 1.79218726 | 1.89611986 | 1.17300152 | 1.22138908 | 1.35025166 |

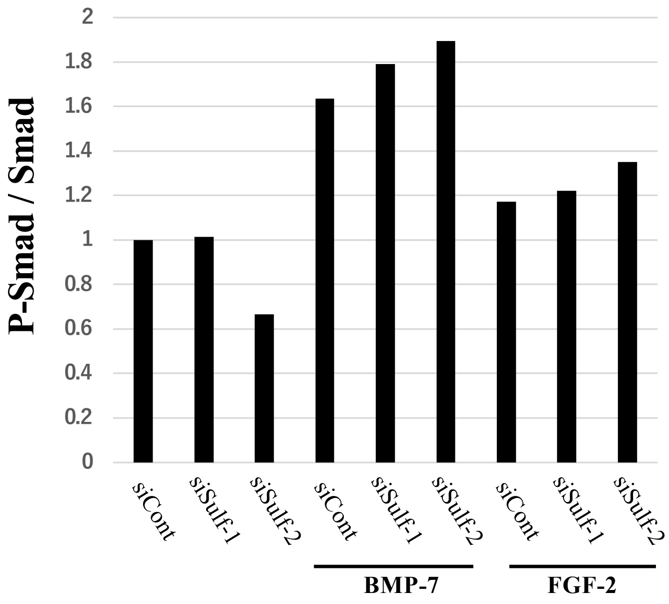

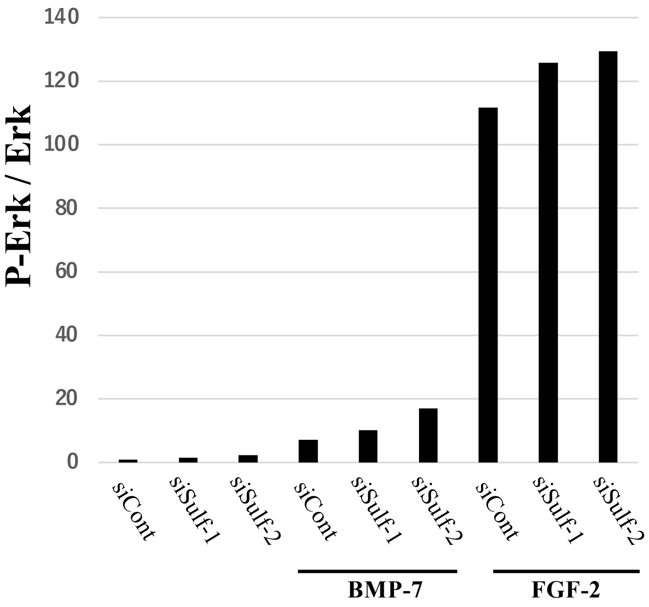

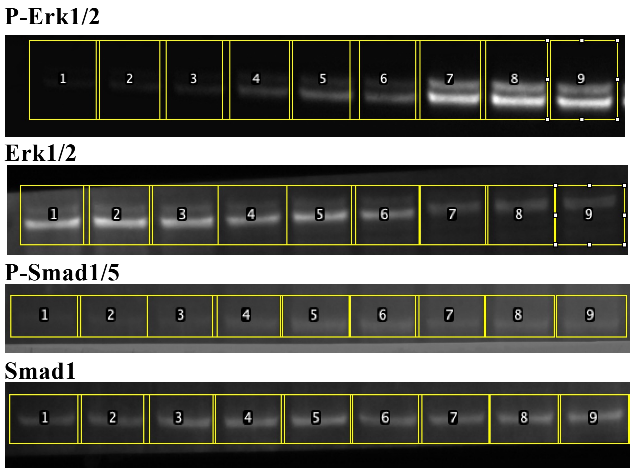

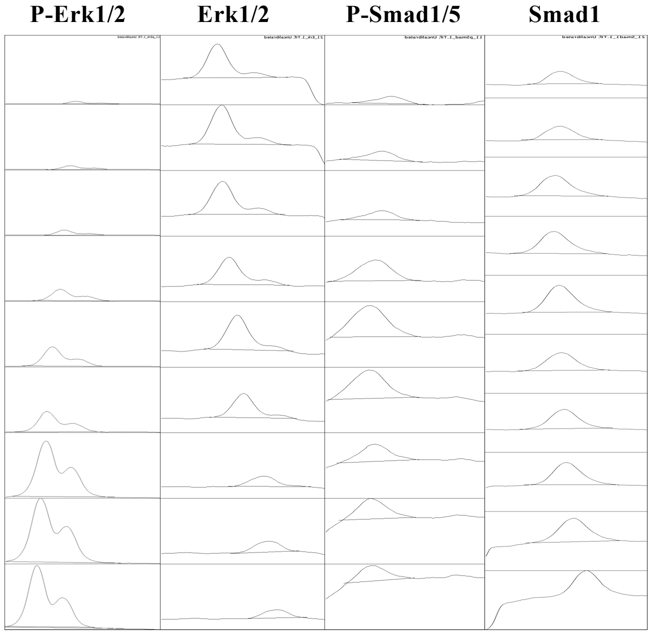

### Suppl. Fig. 4: Densitometry corresponding to Figure 1E

| | Control | | | BMP7 | | | FGF2 | | |
|---|---|---|---|---|---|---|---|---|---|
| **P-Erk/Erk ratio** | **siCont** | **siSulf-1** | **siSulf2** | **siCont** | **siSulf-1** | **siSulf2** | **siCont** | **siSulf-1** | **siSulf2** |
| Sample 1 | 1 | 1.43 | 2.16 | 1.49 | 1.83 | 2.02 | 10.05 | 15.65 | 16.1 |
| Sample 2 | 1 | 0.91 | 0.93 | 1.19 | 1.04 | 1.12 | 3.81 | 4.98 | 5.36 |
| Sample 3 | 1 | 1.09 | 0.68 | 1.12 | 1.24 | 2.3 | 7.17 | 8.76 | 11.93 |
| Sample 4 | 1 | 1.68 | 1.47 | 4.65 | 3.62 | 3.42 | 7.58 | 11.95 | 11.61 |
| Sample 5 | 1 | 1.49 | 2.26 | 7.23 | 10.19 | 17 | 111.7 | 125.8 | 129.4 |
| **Mean** | **1** | **1.32** | **1.5** | **3.136** | **3.584** | **5.172** | **28.062** | **33.428** | **34.88** |
| **SD** | **0** | **0.31288976** | **0.70911917** | **2.72091529** | **3.83025195** | **6.66283123** | **46.8078377** | **51.787419** | **52.9772229** |
| **Student's t-test** | | **0.04207286** | **0.09500307** | | **0.27084141** | **0.13454714** | | **0.04162963** | **0.03638694** |
| **P-Smad/Smad ratio** | | | | | | | | | |
| Sample 1 | 1 | 0.65 | 0.79 | 1.43 | 1.05 | 1.11 | 0.84 | 0.52 | 0.74 |
| Sample 2 | 1 | 1 | 1.03 | 2.6 | 2.19 | 2.14 | 1.42 | 1.41 | 0.99 |
| Sample 3 | 1 | 0.77 | 0.95 | 0.65 | 0.38 | 0.034 | 0.4 | 0.1 | 0.016 |
| Sample 4 | 1 | 0.78 | 0.75 | 2.33 | 1.48 | 1.61 | 1.57 | 1.48 | 1.24 |
| Sample 5 | 1 | 1.01 | 0.67 | 1.64 | 1.79 | 1.9 | 1.17 | 1.22 | 1.35 |
| **Mean** | **1** | **0.842** | **0.838** | **1.73** | **1.378** | **1.3588** | **1.08** | **0.946** | **0.8672** |
| **SD** | **0** | **0.15738488** | **0.14805404** | **0.77158927** | **0.69697202** | **0.83415658** | **0.47005319** | **0.60677838** | **0.53100772** |
| **Student's t-test** | | **0.04407543** | **0.03534825** | | **0.04620204** | **0.04842942** | | **0.05060637** | **0.06692027** |

**Data correspond to Figure 1E**

| | | | | | | | | | | |
|---|---|---|---|---|---|---|---|---|---|---|
| **Sample 1** | **P-Erk/Erk** | 0.3132761 | 0.4481137 | 0.6757126 | 0.4658265 | 0.5742594 | 0.6316884 | 3.149565 | 4.9012778 | 5.0429185 |
| | **P-Smad/Smad** | 0.9414203 | 0.6132228 | 0.7445879 | 1.3455544 | 0.9904497 | 1.0453845 | 0.7892719 | 0.4855086 | 0.6939785 |
| | **Ratio** | | | | | | | | | |
| | **P-Erk/Erk** | 1 | 1.4304111 | 2.1569231 | 1.4869517 | 1.8330773 | 2.0163948 | 10.053638 | 15.645232 | 16.097359 |
| | **P-Smad/Smad** | 1 | 0.6513805 | 0.7909197 | 1.4292812 | 1.0520802 | 1.1104334 | 0.8383841 | 0.5157193 | 0.7371612 |
| **Sample 2** | **P-Erk/Erk** | 0.25601657 | 0.23376099 | 0.23698687 | 0.30528115 | 0.26554968 | 0.28735514 | 0.97600114 | 1.27521127 | 1.37215899 |
| | **P-Smad/Smad** | 0.430745 | 0.43109659 | 0.44210655 | 1.12011308 | 0.94270202 | 0.92081453 | 0.61204027 | 0.60859057 | 0.42666213 |
| | **Ratio** | | | | | | | | | |
| | **P-Erk/Erk** | 1 | 0.91306977 | 0.92567002 | 1.19242729 | 1.03723631 | 1.12240837 | 3.81225769 | 4.98097159 | 5.35964911 |
| | **P-Smad/Smad** | 1 | 1.00081624 | 1.02637651 | 2.60040878 | 2.18853852 | 2.13772541 | 1.42088771 | 1.41287902 | 0.99052138 |
| **Sample 3** | **P-Erk/Erk** | 0.2006922 | 0.21944442 | 0.136508586 | 0.224491246 | 0.248059805 | 0.461954021 | 1.439850606 | 1.758376584 | 2.395222526 |
| | **P-Smad/Smad** | 2.457923317 | 1.894221823 | 2.346990224 | 1.587901338 | 0.939694237 | 0.084173695 | 0.989273556 | 0.252193189 | 0.039399458 |
| | **Ratio** | | | | | | | | | |
| | **P-Erk/Erk** | 1 | 1.093437711 | 0.680188793 | 1.118584805 | 1.236021155 | 2.301803561 | 7.174422346 | 8.761559155 | 11.93480625 |
| | **P-Smad/Smad** | 1 | 0.770659446 | 0.954867146 | 0.646033718 | 0.382312268 | 0.034245859 | 0.40248349 | 0.102604173 | 0.016029572 |
| **Sample 4** | **P-Erk/Erk** | 0.21996965 | 0.36958888 | 0.32299555 | 1.02343835 | 0.79640473 | 0.75255414 | 1.66663707 | 2.62837957 | 2.55483613 |
| | **P-Smad/Smad** | 0.73356315 | 0.57037382 | 0.55138057 | 1.70920961 | 1.0845948 | 1.17794949 | 1.15519592 | 1.08314197 | 0.90982075 |
| | **Ratio** | | | | | | | | | |
| | **P-Erk/Erk** | 1 | 1.68018123 | 1.46836415 | 4.65263434 | 3.62052096 | 3.42117258 | 7.57666827 | 11.9488282 | 11.6144938 |
| | **P-Smad/Smad** | 1 | 0.77753882 | 0.75164705 | 2.33001019 | 1.47852956 | 1.60579153 | 1.57477366 | 1.47654905 | 1.24027597 |
| **Sample 5** | **P-Erk/Erk** | 0.09498524 | 0.14178853 | 0.21424583 | 0.6871443 | 0.96766794 | 1.61479074 | 10.6115642 | 11.9452033 | 12.2929663 |
| | **P-Smad/Smad** | 0.5508664 | 0.55851051 | 0.36730307 | 0.90104139 | 0.98725574 | 1.04450871 | 0.64616712 | 0.6728222 | 0.74380827 |
| | **Ratio** | | | | | | | | | |
| | **P-Erk/Erk** | 1 | 1.49274274 | 2.25556964 | 7.23422155 | 10.1875607 | 17.0004379 | 111.718029 | 125.758516 | 129.419748 |
| | **P-Smad/Smad** | 1 | 1.01387653 | 0.66677343 | 1.63568044 | 1.79218726 | 1.89611986 | 1.17300152 | 1.22138908 | 1.35025166 |

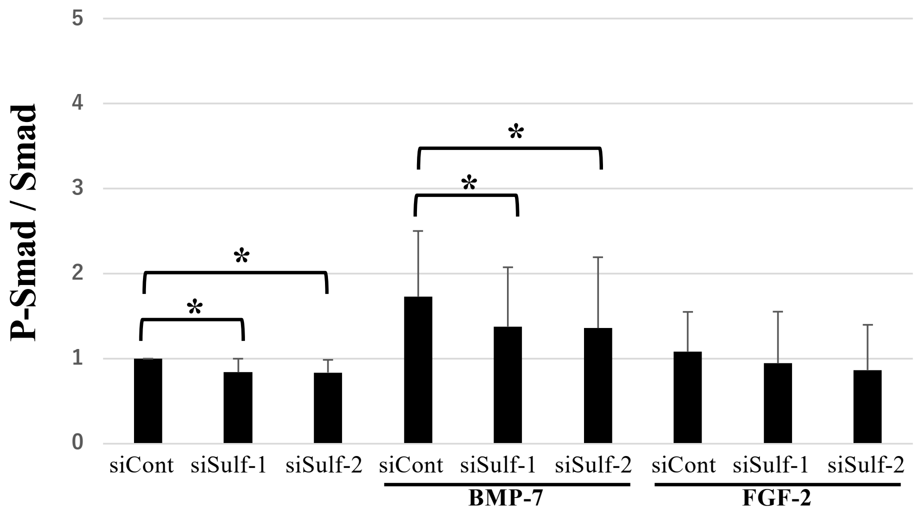

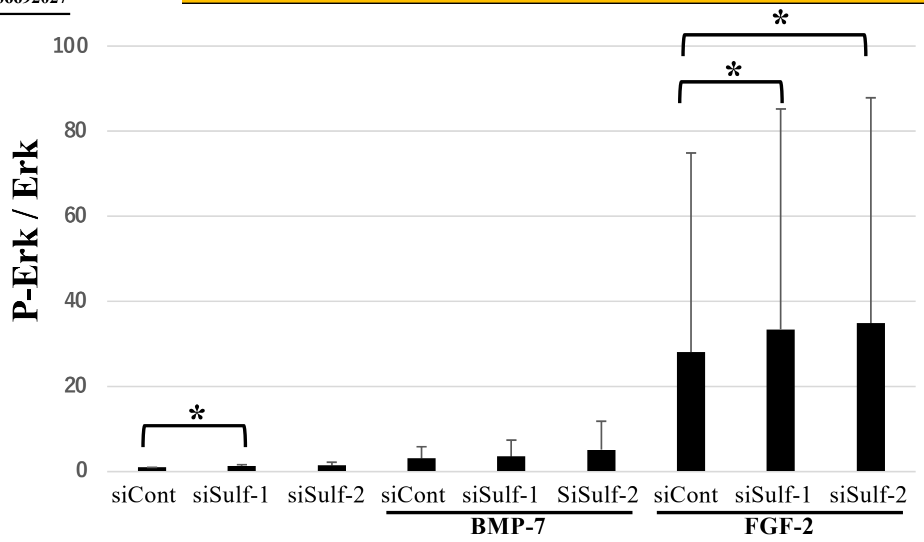
